# SBMLNetwork: a framework for standards-based visualization of biochemical models

**DOI:** 10.1101/2025.05.09.653024

**Authors:** Adel Heydarabadipour, Lucian Smith, Joseph L. Hellerstein, Herbert M. Sauro

## Abstract

SBMLNetwork is an open-source software library that makes the SBML Layout and Render packages practical for standards-based visualization of biochemical models. Current tools often manage model visualization data in custom-designed, tool-specific formats and store it separately from the model itself, hindering interoperability, reproducibility, and the seamless integration of visualization with model data. SBMLNetwork addresses these limitations by building directly on the SBML Layout and Render specifications, automating the generation of standards-compliant visualization data, offering a modular implementation with broad integration support, and providing a robust API tailored to the needs of systems biology researchers. We illustrate the capabilities of SBMLNetwork across key visualization tasks, including SBGN-compliant visualization, application of predefined style templates, layout arrangement to reflect pathway logic, and integration of model data into network diagrams. These examples demonstrate how SBMLNetwork enables high-level visualization features and seamlessly translate user intent into reproducible outputs that support both structural representation and dynamic data visualization within the SBML model. SBMLNetwork is freely available at https://github.com/sys-bio/SBMLNetwork under the MIT license.

## Introduction

Visualization is an essential element in systems biology, particularly for interpreting complex biological models. It transforms multifaceted datasets—ranging from network topologies to simulation results—into clear, intuitive graphical representations that reveal the underlying interactions and dynamic behaviors of the models. By visually depicting biological models, researchers can deepen their understanding of biological systems through the observation of network topology, system dynamics, and biological responses. Beyond enhancing model comprehension, visualization also enables effective collaboration by allowing researchers to clearly communicate their insights and findings to a broader audience.

In recent years, rapid advances in high-throughput technologies have led to an exponential increase in the scale and complexity of biological data, shifting the research emphasis from data acquisition to data interpretation [1]. This increased focus has reemphasized the significance of data visualization and spurred the development of specialized tools for the visualization of biological models. Notable examples include Escher [1], which enables rapid and semi-automated design as well as visualization of biological pathways and associated data [1]; CellDesigner, a tool that facilitates the construction of both gene-regulatory and biochemical networks [2]; MINERVA, a comprehensive platform for curating, annotating, and visualizing molecular interaction networks [3]; CySBML and cySBGN, which support the seamless import and management of SBML (Systems Biology Markup Language) [4] and SBGN (Systems Biological Graphical Notation) [5] formats in Cytoscape [6, 7]; and Newt, designed for creating pathways in SBGN format [8]. However, the concurrent development of the tools has introduced various data formats for storing visualization data, leading to challenges such as limited interoperability, cumbersome data exchange, and difficulties in reproducing model visuals across different platforms [9]. The emergence of these challenges underscores the importance of developing widely adopted standards for biological model visualization.

Mirroring the role of formats like SBML for storing and exchanging model data, visualization standards should aim to ensure that all generated data remain seamlessly exchangeable among different tools, reproducible in future analyses, and extensible for further developments. A robust standard for storing visualization data must address both the arrangement and stylistic properties of biological model elements, ensuring that any tool can interpret them without relying on rigid, tool-specific definitions. Moreover, a key requirement for each visualization standard should be the explicit mapping of model components to their corresponding visual elements. This mapping enables straightforward cross-referencing between each model visual and its underlying model entity, a critical step in seamlessly integrating simulation and experimental data with the graphical representation of the model. In addition, the standard must facilitate unified storage of data, enabling both model and visualization data to be maintained within a single file. This simplifies model management by eliminating the need to manage multiple files and reducing errors when linking corresponding elements to their graphical representations.

Among the candidate standards for storing visualization data, SBML with Layout [10] and Render [11] packages stands alone in offering robust, tool-agnostic support for both the positioning and graphical styling of model elements. It also provides explicit mapping between each visual element and its corresponding model component. In addition, it offers the functionality to store visualization data in the same file as the SBML model data. However, despite its significant potential to serve as the de facto standard for model visualization, SBML with Layout and Render packages has remained underutilized. Its limited adoption can largely be attributed to the fact that the process for generating and editing visualization data for a biological model using this standard is very tedious—mainly because its steep learning curve demands a significant investment of time and technical expertise, and its API is not particularly intuitive for non-technical users. SBMLDiagrams [12] was an important early effort to address some of these limitations by providing a higher-level Python interface to the Layout and Render specifications. However, its reliance on manual placement of visual elements and its lightweight wrapper architecture meant that many of the core usability challenges remained unresolved. Consequently, these barriers have continued to limit the widespread adoption of SBML Layout and Render and have prevented the standard from reaching its full potential in the field.

In this paper, we present SBMLNetwork—a software tool designed to leverage the unique potential of SBML with Layout and Render packages and promote its adoption as the standard for biological model visualization. SBMLNetwork addresses current limitations of SBML with Layout and Render by incorporating an auto-layout algorithm that efficiently generates initial visualization data, along with a comprehensive yet user-friendly API that enables intuitive customization of the detailed features of model visuals. In addition, as a modular command-line tool that integrates easily with other projects, SBMLNetwork facilitates seamless adoption across diverse research workflows and software environments.

## Design and Implementation

SBMLNetwork is built on a modular, layered architecture that organizes functionalities into discrete, interrelated levels. This design—illustrated in Figure 1—enables us to isolate key responsibilities such as standard compliance, input/output data management, core processing, cross-language integration, and user interaction, so that each module can be developed, maintained, and enhanced independently. This separation promotes robustness, scalability, and seamless interoperability across diverse platforms. In the following sections, we detail each layer, outlining its specific role, the technologies employed, and its interactions with adjacent layers.

**Fig 1.**
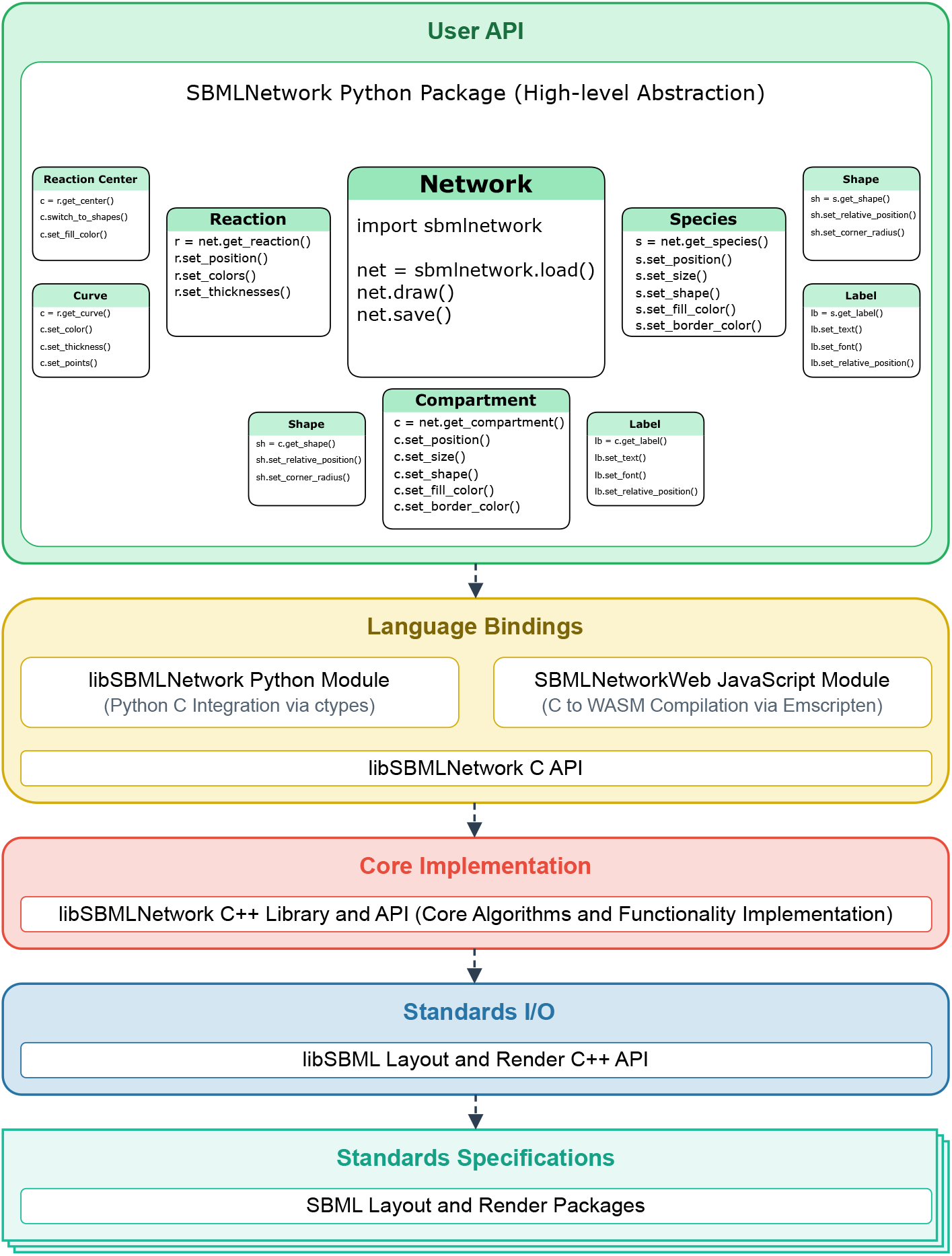
SBMLNetwork Layered Architecture. The figure illustrates the modular, multi-layered design of SBMLNetwork, which is organized into discrete levels responsible for standard compliance, I/O operations, core processing, cross-language integration, and user interaction. In particular, the User API Layer is outlined with its three sub-layers that offer a seamless progression from high-level operations to granular control over visualization details and provide support for both novice and expert users.

### Standards Specification Layer

At the core of our architectural design, is the SBML Layout and Render packages that provides a standardized visualization of reaction networks, expressed in biochemical terms. The SBML Layout package comprises specific layout elements that mirror the structure of the core SBML model and record the positions and sizes of its elements. It also allows a single model element to be represented by multiple layout elements (alias elements) [13, 14] and supports cases where a layout element does not directly correspond to any model element, for example when depicting sources and sinks in a biological reaction. Complementary to the Layout package, the SBML Render package independently manages visual styling, such as colors, node shapes, line styles, and fonts, by organizing a set of styles that can be applied to layout elements. The style attributes in these sets are primarily derived from corresponding elements in the SVG specification [11]. Using the SBML Layout and Render packages, our system reliably supports both the positioning and visual styling of model elements, explicitly links each visual element with its corresponding model component, and stores visualization data together with the core SBML model data.

### Standards I/O Layer

Building on the foundation provided by the SBML Layout and Render packages, this layer ensures that our system’s input and output operations strictly adhere to the standards defined by these packages. To achieve this, we leverage libSBML [15]—a mature, high-level API library with built-in support for the Layout and Render packages—to efficiently read, write, manipulate, and validate SBML Layout and Render data. This approach guarantees that our system seamlessly and reliably manages the visualization of biological models according to these established standards.

### Core Implementation Layer

At the heart of our system lies the Core Implementation Layer, which encompasses the fundamental algorithms and functionalities required for the automated generation and management of SBML Layout and Render elements. This layer offers several key capabilities that work together to ensure that visualization data for each model element is generated and managed with high accuracy and reliability. Examples of these capabilities include: the automatic generation of Layout and Render elements for each model element in SBML models that lack such data; the creation of alias elements when a single model element requires multiple visual representations [13]; the automatic assignment of default or element-specific styles to each model element; providing users with access to a set of HTML colors (with corresponding hex codes) [16] and harmonic stylistic features for setting a consistent theme across the network; the automatic initial arrangement of visual elements through a refined force-directed autolayout: an adaptation of the Fruchterman–Reingold algorithm [17] augmented with Bézier-curve alignment and expressly tuned for biological network models [18]; and a robust error logging system that captures function failures and provides diagnostic feedback, guiding users toward identifying and resolving issues encountered during operation.

### Language Bindings Layer

This layer bridges the gap between the underlying C++ implementation and higher-level applications by exposing core functionalities via a robust C API and native interfaces. The provided C API abstracts the complexities of the core implementation and enables seamless integration with multiple programming languages, ensuring that the main capabilities of our system are reliably accessible regardless of the language used. Then, by leveraging the provided C API, we make our system accessible to various programming environments through native interfaces. For example, to offer our users a simple interface that facilitates incorporation into scientific computing and data analysis workflows, our Python bindings—developed using ctypes [19]—integrate the core C API with Python applications. Additionally, to meet the rising demand for web-based applications, we provide JavaScript bindings by compiling the core C/C++ source code into WebAssembly (WASM) [20] using Emscripten [21], which delivers our core functionality directly in the browser and enables seamless integration into web-based applications. This design approach not only promotes reusability but also simplifies the development and maintenance of language-specific bindings and further enhances our system’s extensibility.

### User API Layer

Our User API Layer exposes the core functionalities of our system to end users by leveraging the Python bindings from the Language Bindings Layer and abstracting its intrinsic complexity into an intuitive API. Its design approach targets systems biology researchers as the primary users and aims to significantly reduce the learning curve for them to start using our system. As shown in Figure 1, this layer is organized into three sub-layers, each offering a different level of abstraction.

At the top-most sub-layer, called Network, users can perform essential operations such as loading an SBML model, rendering its figure, and saving the model along with added or modified visualization data. This sub-layer also provides highly abstracted visualization features critical for model analysis and interpretation, including seamless data integration, dynamic grouping of model elements based on stylistic attributes, and batch application of modifications across multiple elements.

The middle sub-layer is designed for power users who require direct access to the visual representations of the biological model’s components, including Species, Reactions, and Compartments. With this layer, users can directly access network components and manipulate both their arrangement and stylistic attributes.

Finally, for expert users, the lowest sub-layer offers direct access to the graphical building blocks that constitute the model’s visual representation, such as geometric shapes, curves, and labels. This granular level of control is essential for users who need to rigorously customize or extend the default visualization and manipulate every detail of the rendered network.

Additionally, this layer integrates skia-python [22]— a robust Python binding for the Skia 2D rendering engine—to deliver high-quality outputs of the visualized model in multiple formats, including JPEG, PNG, SVG, and PDF.

Overall, our layered architecture delivers a robust and flexible system for generating and managing visualization data for biological models by leveraging the SBML Layout and Render standards. Performance-critical operations are implemented in C++, while tools such as ctypes and Emscripten enable seamless interaction across diverse programming environments and enhance cross-platform compatibility. At the forefront of this architecture, the User API Layer plays a pivotal role. It is a carefully structured interface that abstracts the underlying complexity into an accessible, multi-tiered framework. By offering a seamless progression from high-level operations to granular control over model elements and visualization details, the User API Layer meets the specific needs of both novice and expert users. Furthermore, the system’s modular design allows each layer to be updated or replaced independently, which facilitates the integration of new language bindings without impacting core functionalities. Given the evolving nature of visualization requirements, this approach not only addresses current needs but also provides a solid foundation for future enhancements, ensuring that the system evolves alongside emerging technologies and user demands.

## Results

In this section, we demonstrate how SBMLNetwork addresses a variety of high-demand visualization tasks for biological models by providing seamless support for SBGN-compliant representations, advanced styling capabilities, flexible network element arrangements, and direct data integration on top of the visualized network—all while preserving arrangement and styling information within the SBML file.

### Visualization of SBGN-Compliant Networks

SBGN [5] is a widely adopted standard for visualizing biological models, enabling the unambiguous representation of complex biological knowledge through a set of easily recognizable glyphs. By providing the following visualization features, SBMLNetwork supports the generation of SBGN-compliant visualizations: (i) support for multi-compartment models; (ii) the ability to assign multiple shapes and labels to each network element; (iii) support for visualizing reaction centers as distinct geometric shapes; (iv) the capability to render multi-segmented curves for reaction paths; (v) support for demonstrating empty species; and (vi) the flexibility to use various shapes for arrowheads on reaction curves. To demonstrate this capability, we recreated a well-known SBGN-compliant network, the iNOS pathway example [23]. The recreated figure, shown in Figure 2, demonstrates that SBMLNetwork can faithfully and efficiently produce high-quality SBGN visualizations. This capability enables modelers to fully leverage the advantages of the SBGN stylistic format while storing visualization data alongside the core SBML model.

**Fig 2.**
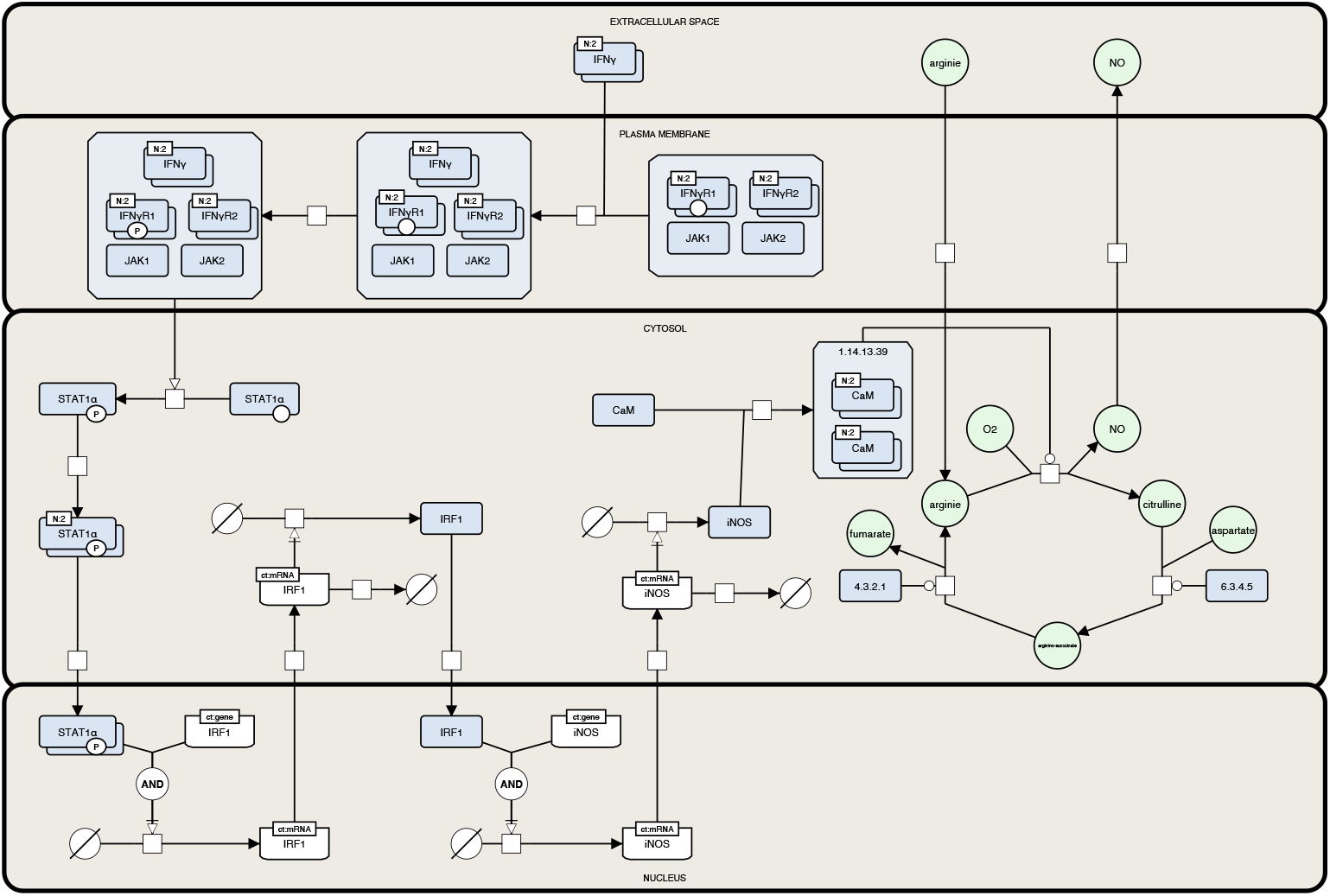
Recreated iNOS Pathway Example in SBGN Format. This figure shows the iNOS pathway example [23] recreated using SBMLNetwork. The visualization adheres to the SBGN standard, demonstrating the tool’s ability to faithfully and efficiently generate high-quality visualizations of complex biological networks while storing visualization data alongside the core SBML model.

### Visual Styling

Visual styling is an effective technique for achieving aesthetically consistent and insightful visualizations of biological models. SBMLNetwork, with its comprehensive customization capabilities, enables users to apply a variety of styles to their models. To do so, SBMLNetwork provides a set of predefined style templates that users can select to alter the overall appearance of their network visualizations. Among these style templates is Escher [1]—a widely recognized scheme for depicting metabolic networks. The capability of our tool to apply the Escher style template is especially advantageous for users because the visualization data produced by tools that export to the Escher format omits explicit styling information. With support for the following features: (i) full control over the color and thickness of reaction curves; (ii) customization of font colors and sizes for network element labels; (iii) flexible configuration of multiple shapes for reaction centers; and (iv) configurable settings for arrowhead designs on reaction curves, SBMLNetwork is capable of setting the Escher style template on biological models stored in SBML format. To demonstrate this capability, as shown in Figure 3, we applied the Escher style to the core *E. coli* metabolic model [24], where we first generated the SBML model with its Layout data using EscherConvertor [25], and then used SBMLNetwork to create Render data and apply the Escher style template on it.

**Fig 3.**
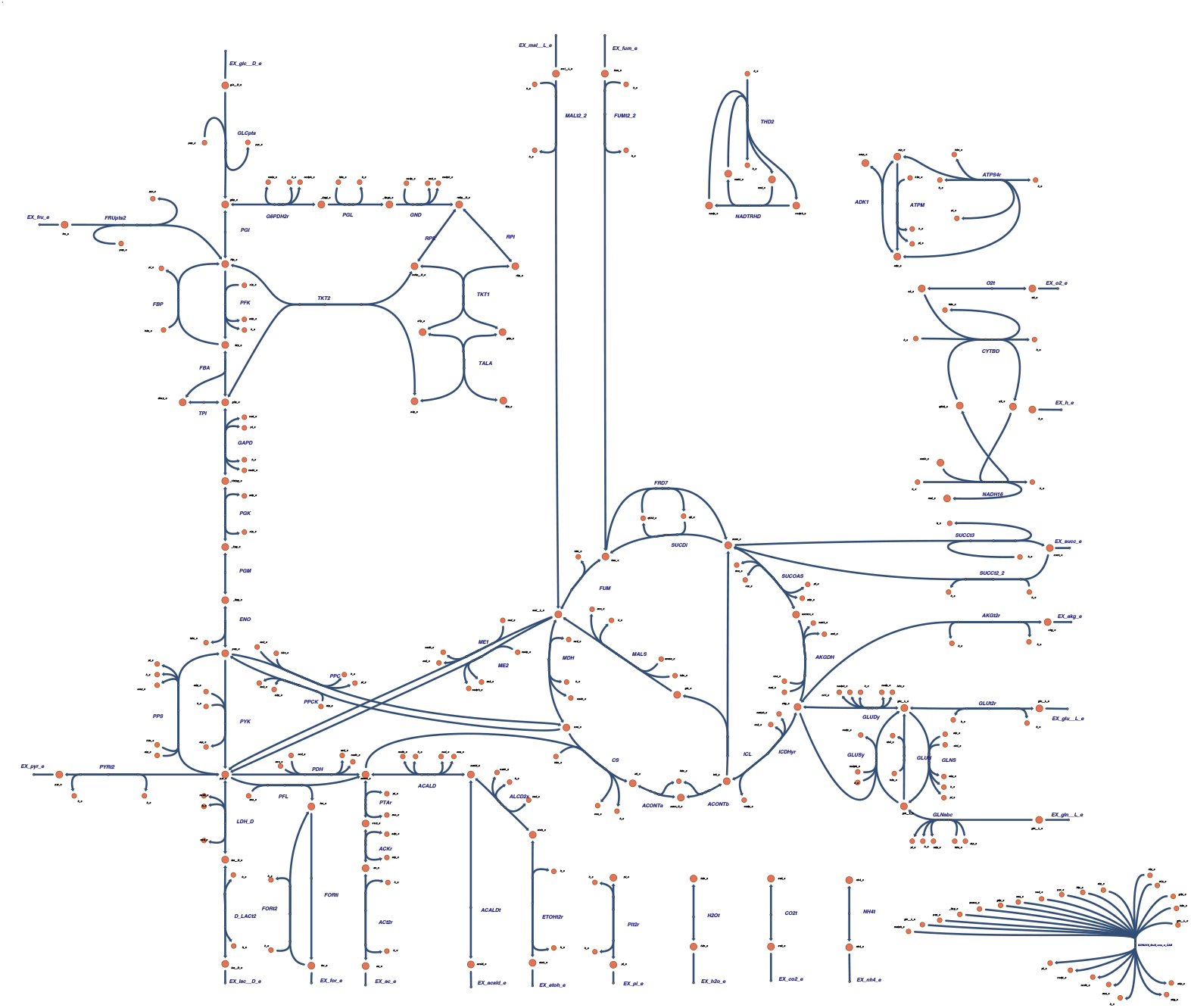
Applying Escher-Style Template on the Core E. coli Metabolic Model. This figure demonstrates the capability of SBMLNetwork to apply a style template to a complex biological model. The core *E. coli* metabolic model [24] was chosen, and its SBML Layout data was generated using EscherConvertor. Subsequently, SBMLNetwork was employed to create SBML Render data and apply the well-known Escher style on the model. The resulting visualization showcases the advanced styling capabilities of SBMLNetwork, ensuring a uniform and fine-grained representation of complex biological networks in specific styles.

The resulting visualization demonstrates that SBMLNetwork effectively supports advanced visual styling for complex biological models, enabling modelers to achieve highly customized visualizations and maintain uniform styling across the entire model down to its finest details.

### Network Arrangement

A well-organized layout is essential for conveying the logic of reactions in a biological model. By carefully arranging network elements, users can highlight relationships, directional flows, and hierarchical structures. Inspired by widely used pathway maps such as KEGG Metabolic Pathways [26] and Escher pathway maps [1], SBMLNetwork provides an advanced alignment feature that positions reactions horizontally, vertically, or along circular arcs.

We demonstrated this capability by visualizing the tricarboxylic acid (TCA) cycle. The Python script in Listing 1 is used to arrange the core TCA reactions in a circular layout and align the pyruvate dehydrogenase step vertically above the cycle. The resulting visualization, shown in Fig. 4, highlights metabolite flow and the pathway’s cyclic nature. Notably, this precise arrangement is achieved with only a handful of SBMLNetwork commands, demonstrating that even complex positioning tasks can be carried out with minimal coding effort using the SBMLNetwork API. Moreover, because the alignment functionality can be applied to arbitrary subsets of reactions, it scales naturally to larger networks, enabling users to orient pathways and expose underlying biochemical logic with minimal effort.

**Fig 4.**
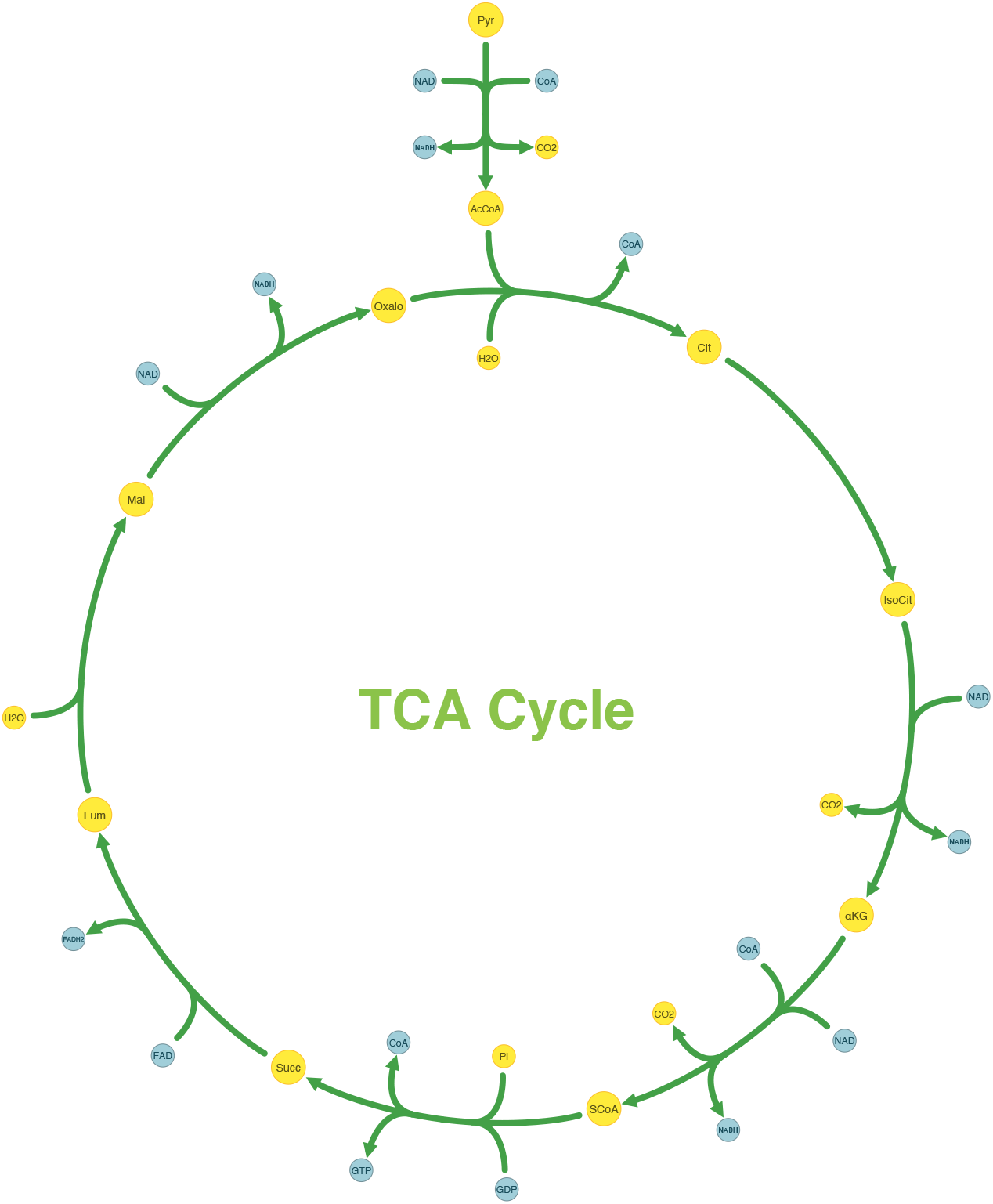
Reaction Alignment of the Tricarboxylic Acid (TCA) Cycle Metabolic Network Model. Generated using the Python script in Listing 1, SBMLNetwork’s arrangement feature positioned the pyruvate dehydrogenase reaction vertically at the top and arranged the core TCA-cycle reactions in a circular layout to emphasize metabolite flow and the pathway’s cyclic nature. Metabolites are shown in yellow, cofactors in blue, and reactions in green to distinguish the different components of the pathway.

### Data Integration

Effective visualization not only illustrates the structure of biological models but also integrates model data, enabling researchers to explore both network topology and dynamic behaviors. SBMLNetwork supports this integration by overlaying dynamic values—such as reaction fluxes and species concentrations—onto the visualized network through color-coded and size-coded mappings, providing an intuitive view of metabolic activity over time. To ensure accurate and accessible displays, SBMLNetwork offers user-friendly helper functions, a fully customizable color bar for gradient encodings, and size-scaling utilities that map model data to network elements. Importantly, all visual elements related to data integration are seamlessly preserved alongside the model within the SBML Layout and Render specifications.

To demonstrate this feature, we used SBMLNetwork to render reaction flux data for the glycolysis pathway in a model of Saccharomyces cerevisiae [28]. Simulation data was provided by the Tellurium package [29]. Fluxes at a specific simulation time point were mapped to a gradient color scale—vivid hues for high flux and muted tones for low—and applied to the reaction curves, yielding the clear snapshot shown in Figure 5. SBMLNetwork offers the same workflow for mapping species concentrations to the fill color of species nodes in the network. In addition, it can encode reaction fluxes in the thickness of reaction curves and species concentrations in the size of species nodes. By integrating both reaction-flux and species-concentration data into network visualizations, SBMLNetwork provides a robust platform for dynamic exploration of biological models, ensuring that structural context and dynamic behavior are conveyed cohesively and with clear visual emphasis.

**Listing 1.**
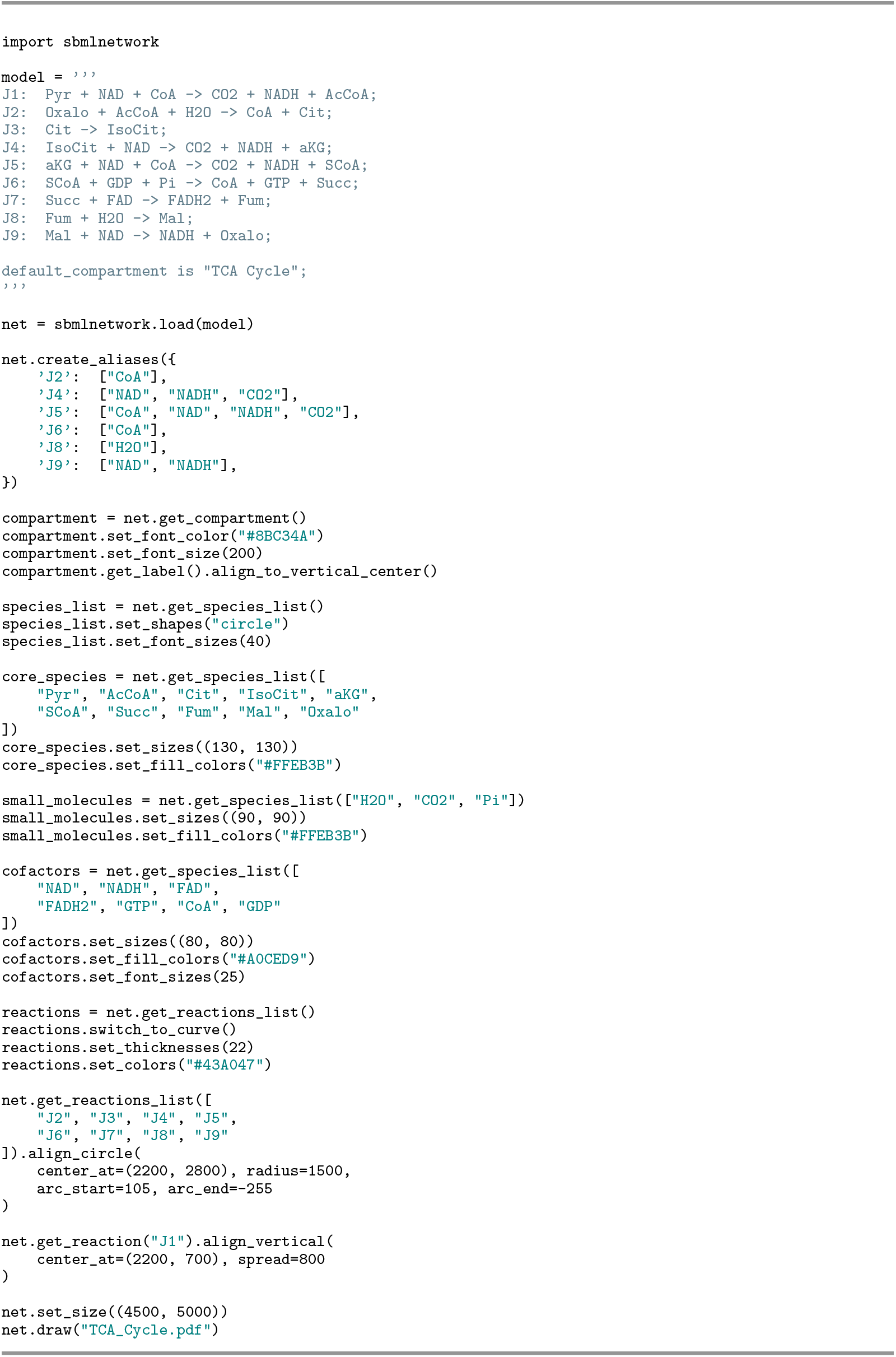
SBMLNetwork Python script that loads the TCA-cycle model written in Antimony [27] format, creates reaction-specific alias species, applies customized styling to compartments, species, and reactions, aligns the pyruvate dehydrogenase reaction vertically at the top, and arranges the main cycle reactions in a circular layout to generate the visualization shown in Fig. 4.

**Fig 5.**
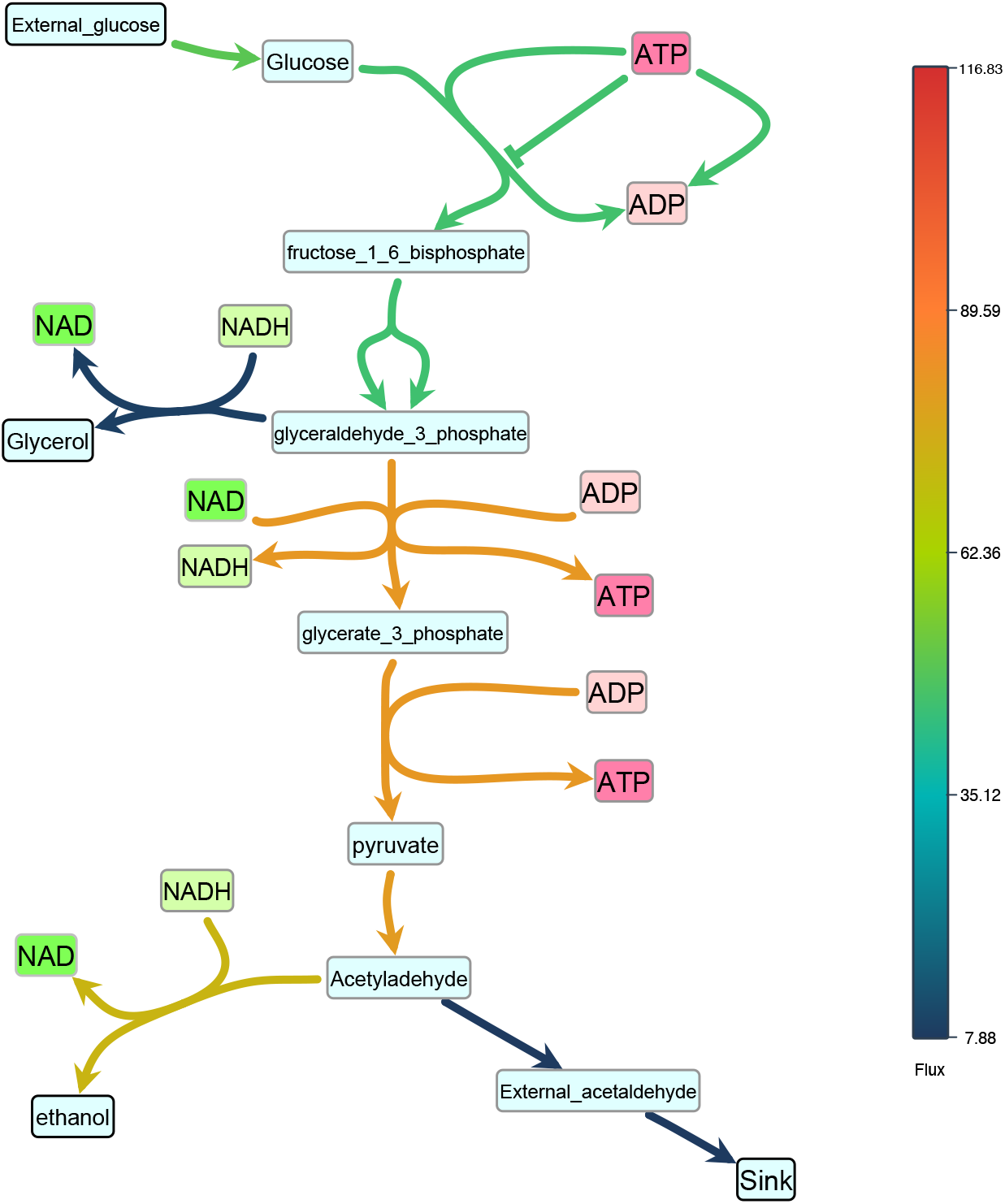
Color-encoded reaction fluxes in the *S. cerevisiae* glycolysis pathway. SBMLNetwork maps reaction-flux values from the glycolysis pathway in a model of Saccharomyces cerevisiae [28] to a continuous color gradient—vivid hues for high fluxes and muted tones for low, which offers a visual snapshot of pathway activity at a selected simulation time point.

## Discussion

In this work, we introduced SBMLNetwork, a software tool that streamlines and strengthens the visualization of complex biological models. By embracing SBML’s Layout and Render standards—and removing the steep technical hurdles that have hindered their wider adoption—SBMLNetwork directly addresses three persistent challenges: (i) poor interoperability across current visualization tools, (ii) visualization formats that cannot be seamlessly linked to their underlying model data, and (iii) the lack of a unified way to store model and visualization information together in a single file.

Building on this foundation, SBMLNetwork is architected as a multi-layered system. In the lower layers, performance-critical operations are implemented in C++ and ensure strict adherence to SBML Layout and Render standards. The intermediate layers facilitate adaptation and extension through language-specific bindings, with support provided for both Python and JavaScript. At its highest level, the user-facing APIs are tailored for non-technical users and high-demand use cases where an intuitive interface to the underlying functionalities is provided for end users.

To demonstrate SBMLNetwork’s capability to address key visualization challenges in biological modeling, we applied the tool to several common use cases. For instance, our recreated iNOS pathway demonstrates SBMLNetwork’s ability to generate visualizations that adhere to the SBGN standard—a widely adopted approach for depicting biological models—while preserving every detail of the model’s layout and styling information. In another example, applying an Escher-style template to a model underscores that SBMLNetwork’s advanced styling features can be leveraged to achieve the desired visual format with minimal effort, effectively compensating for the lack of built-in styling options in the available tools. Additionally, the network arrangement features—exemplified by the visualization of the TCA Cycle metabolic network—eliminate the tedious task of manually positioning reactions by aligning them in orientations that reveal the underlying biochemical logic with minimal coding effort. Finally, by overlaying simulation data—such as reaction fluxes encoded with gradient color mappings—onto network visualizations and including informative elements such as a fully customizable color bar, SBMLNetwork seamlessly integrates a model’s structure with its dynamic behavior. Collectively, these examples demonstrate that SBMLNetwork meets high-demand visualization requirements and provides a robust platform for exploring and interpreting complex biological models.

In summary, SBMLNetwork addresses critical visualization challenges in systems biology while unifying structural and stylistic data in a single file alongside the core SBML model. By lowering the technical barriers associated with using the SBML Layout and Render packages and providing a variety of user-focused functionalities, our tool facilitates reproducible, interoperable, and seamless model visualizations. Furthermore, SBMLNetwork lays a solid foundation for future developments in standards-based biological model visualization—an increasingly important component of large-scale, data-driven research in systems biology—owing to its highly extensible design that enables seamless integration with a wide range of tools and workflows.

## Availability and Future Directions

SBMLNetwork is maintained as an open-source project on GitHub (https://github.com/sys-bio/SBMLNetwork), and is distributed under the MIT License. Detailed build instructions for compiling the binaries from source are available for developers on the GitHub page. The release page provides both static and dynamic binary distributions of the underlying C/C++ libraries, as well as the WebAssembly and JavaScript bindings. Additionally, SBMLNetwork is distributed as a Python package and can be installed from PyPI using the command pip install sbmlnetwork. The source code used to generate Figures 2–5, along with the output SBML file containing the visualization data in SBML Layout and Render format, is also available on the GitHub repository.

As we look ahead to the future of SBMLNetwork, its immense potential—built primarily upon its robust technical foundation—sets the stage for next-generation advancements in biological model visualization. This strong foundation is reflected in its robust C++ backend, which provides the high-performance computations required for processing complex models; its multi-layered architecture, which offers outstanding modularity and scalability; and its integrated high-level language bindings, which facilitate seamless integration into a variety of workflows. Leveraging these foundational strengths, the following outline the prospective directions for further research:

- **High-Performance Visualization for Large-Scale Biological Models:** One promising future direction is to fully exploit SBMLNetwork’s robust C++ backend for handling large-scale models. Its high-performance computational capacity offers a unique opportunity to efficiently visualize extensive models—such as those available in the BiGG Models database [30]—that commonly push other available tools to their limits. This direction should be actively pursued to leverage SBMLNetwork’s powerful capabilities and establish it as a premier solution for efficiently scaling the visualization of complex biological models without compromising performance.
- **Next-Generation Simulation Data Integration:** Another promising future direction is to explore innovative methods for overlaying simulation data onto visualized models in SBMLNetwork. For example, recent advances in high-throughput technologies have led to a shift toward representing data as distributions rather than single values. Implementing this approach would enable more effective analysis of multi-omics datasets—capturing variations across different experimental conditions—and result in visualizations that are richer in detail and context [31]. Leveraging its user-friendly API, which provides seamless access to the intricate details of the network, along with its extensible design, SBMLNetwork offers substantial potential as a platform for exploring novel techniques for integrating simulation data onto the visualized model.
- **Rendering Gene Regulatory Networks**. Although SBMLNetwork is primarily geared towards displaying reaction networks, the Render standard is flexible enough to render other types of networks. We have already seen its use to render SBGN networks; however, for the future, one area that is currently poorly served, is rendering gene regulatory networks in a reproducible manner. One of the more useful styles for rendering gene regulatory networks is that generated by BioTapestry [32, 33], and future work could develop a specific BioTapestry style.
- **Predefined Element Styles and Publication-Ready Templates**: Building on SBMLNetwork’s demonstrated capabilities for visual styling, another compelling direction for future developments is to further expand its styling capabilities. One future enhancement involves creating predefined visual styles for individual network elements which allow users to employ more detailed and specialized representations with minimum effort rather than relying solely on basic geometric shapes. For instance, preconfigured skeletal formula depictions could be developed to illustrate carbon-based compounds, or off-the-shelf SBGN-compatible styles could be established for each type of network component. Another styling opportunity is to develop customized visualization templates for the entire network that conform to the formatting standards used by scientific journals. Such templates would significantly facilitate the creation of high-quality, publication-ready figures that are both easy to generate and highly reproducible.
- **AI-Driven Visualization Automation**: SBMLNetwork’s fully scriptable API offers programmatic control over every layout and styling operation, making it an ideal platform for AI-driven visualization automation. Generative AI systems can implement complete visualization pipelines using SBMLNetwork which will produce fully reproducible model visualizations in SBML Layout and Render formats. In addition, machine-learning models trained on curated pathway maps—such as KEGG Metabolic Pathways [26]—can deliver data-driven layout and styling recommendations through the same interface. Together, these capabilities position SBMLNetwork at the forefront of AI-assisted biological network visualization tools, and we anticipate that its growing adoption inspires the community to develop increasingly adaptable, AI-powered workflows in the years ahead.

## Acknowledgments

The authors would like to acknowledge the support of the ‘The Center for Reproducible Biomedical Modeling’, which is funded by the NIH award P41EB023912. In addition, HMS and AH would also like to acknowledge the generous support from NIH award U24EB028887 that funded much of the early work. AH also acknowledges the contributions of Margaret Cook, Stella Anastasakis, Diego Alba Burbano, and Janis Shin whose thorough testing of pre-release versions and thoughtful feedback significantly enhanced the robustness and reliability of this work.

